# Mitochondrial Genome Analysis and Phylogeny and Divergence Time Evaluation of the *Strix aluco*

**DOI:** 10.1101/2023.01.20.524943

**Authors:** Yeying Wang, Haofeng Zhan, Yu Zhang, Zhengmin Long, Xiaofei Yang

## Abstract

In this study, a complete mitochondrial genome of the *Strix aluco* was reported for the first time, with a total length of 18,632 bp. There were 37 genes, including 22 tRNAs, 2 rRNAs, 13 protein-coding genes (PCGs), and 2 non-coding control regions (D-loop). The second-generation sequencing of the complete mitochondrial genome of the *S. aluco* was conducted using the Illumina platform, and then Tytoninae was used as the out-group, PhyloSuite software was applied to build the ML-tree and BI-tree of the Strigiformes, and finally, the divergence time tree was constructed using Beast2.6.7 software, the age of *Miosurnia diurna* fossil-bearing sediments (6.0-9.5 Ma) was set as the internal correction point. The common ancestor of the *Strix* was confirmed to have diverged during the Pleistocene(2.59~0.01Ma). The dramatic uplift of the Qinling Mountains in the Middle Pleistocene and the climate oscillation of the Pleistocene together caused *Strix* divergence between the northern and southern parts of mainland China. The isolation of glacial-interglacial rotation and glacier refuge was the main reason for the divergence of the common ancestor of the *Strix uralensis* and the *S. aluco* during this period. This study provides a reference for the evolution history of the *Strix*.

**Summary statement:** This study was the first time to assemble the complete mitochondrial genome of *Strix aluco*, and report the divergence time of *Strix*. A full discussion was made, and it was inferred that the uplift of the Qinling Mountains and the glacial refuge led to the differentiation of this genus.

## Introduction

*Strix aluco* belongs to Strigiformes, Strigidae, and is a medium-sized owl (Grytsyshina et al., 2016). It is a non-migratory and territorial nocturnal bird (Doña et al., 2016; Sunde et al., 2003). It is widely distributed in the mountain broadleaf forest and mixed forest in Eurasia, and Israel is the southernmost country in the northern hemisphere (Obuch, 2011; Comay et al., 2022). Mammals, fish, amphibians, and even small birds such as sparrows can be its food (Obuch, 2011), and voles are its most preferred food (Solonen et al., 2002; Karell et al., 2009). This species was listed as “Least Concern” (LC) on the IUCN, and the current population trend is stable, the number of said individuals ranges from 1000000 to 2999999. The IUCN (2016). (https://www.iucnredlist.org/), In China, it has been listed as a national class II protected animal.

Mitochondria are characterized by maternal inheritance, high conservation, multiple copies in cells, low sequence recombination rate, and high evolutionary rate, widely used in phylogenetic studies (Yan et al., 2017; Sun et al., 2020), and be able to accurately infer phylogenetic relationships in birds (Tuinen et al., 2000), while complete mitochondrial genomes generally have higher accuracy than partial mitochondrial genes (Haring et al., 2001; Harrison et al., 2004). Previous studies have defined the phylogenetic position of *S. aluco* using a single gene or a combination of multiple mitochondrial genes (Heidrich and Wink, 1994; Yu et al., 2021; Li et al., 2022; Fuchs et al., 2008; Wood et al., 2017). Earlier studies identified the monophyly of Strigiformes phylogeny through the cytochrome B (Cyt B) gene (Wink and Heidrich, 2000), through skeletal comparison, Striginae were divided into three subfamilies: Striginae (13 genera), Surniinae (8 genera) and Asioninae (2 genera) (Salter et al., 2020). Phylogenetic relationships through the cytochrome B (Cyt B) gene also show that the Strigiformes can be divided into four parts, Tytoninae consists of Tytoninae (with *Tyto*) and Phodilinae (with *Phodilus*), Striginae can be divided into Striginae, Surniinae and Ninoxinae. Among them, Striginae consists of six branches (Bubonini+Strigini+Pulsatrigini+Megascopini+Asionini+Otini), and Surniinae consists of three parts: Surnini+Athenini+Aegolini. Ninoxinae is mainly composed of *Ninox*, possibly including *Sceloglaux* (Wink et al., 2009); Zhang et al. (2016) completed the whole mitochondrial genome sequencing of *Asio flammeus* and determined the parallel phylogenetic relationship among the three genera of *Otus, Ptilopsis* and *Asio;* Kang et al. (2018) completed the whole mitochondrial genome sequencing of *Strix uralensis*, and determined the inter-genus relationship of *Otus*+ (*Asio*+ (*Strix*+*Bubo*)) through the study of the mitochondrial genome of Strigidae. Uva et al. (2018) clarified the global distribution of Tytonidae and the time of divergence, and their analysis showed that Tytonidae and *S. aluco* split from a common ancestor dating back to about 45 million years ago. Koparde et al. (2018) identified The juxtaposition phylogenetic relationship between Striginae and Surniinae in the South Asian Subcontinent population with Tytonidae as the out-group. Their study showed that Strigidae and Tytonidae diverged at about 42.5-47.7Ma (mega-annum, million years). The timing of the divergence of the *Strix* is unclear.

There are many reasons for species divergence, among which geological and climatic influences on species diversification cannot be ignored (Claramunt and Cracraft 2015). The Cretacean-Tertiary extinction event was a mass extinction event in Earth’s history that occurred 65 million years ago and wiped out most of the animals and plants of the time, including the dinosaurs. It also wiped out the direct ancestors of tree-dwelling, water birds on Earth today, The few that survived evolved rapidly thereafter (Field et al., 2018). Bird ancestry began to increase exponentially at the end of the Eocene, from 100 species to 10,000 today (Ksepka and Phillips 2015). Since the late Miocene, many birds in the Palaearctic have been migrating on a large scale and changing ranges have led to gene flows that have provided opportunities for the origin of various bird subfamilies (Drovetski 2003; Holm et al., 2014).Climatic oscillation during the Quaternary Period, especially throughout the Pleistocene (2.59~0.01Ma), promoted the evolution of species on a global scale (Hewitt 2000, 2004; Lamb et al., 2019), Pleistocene glacial gyre played a positive role in speciation (Kozma et al., 2018; Zhao et al., 2013; Hung et al., 2014; Mays Jr et al., 2018) and Brito (2005) studied 14 populations of *S. aluco* in Western Europe and found that *S. aluco* in Europe could be divided into three branches originating from three glacial sanctuaries in the Iberian Peninsula, Italy and the Balkan Peninsula. This finding supports the “glacier refuge hypothesis” to describe the origin of S. aluco in Western Europe. The origin and divergence of S. aluco in mainland China are still a mystery.

Divergence time analysis can provide a reference for the evolution process of species and is also the basis for other further studies. In order to clarify the divergence time of species, it is necessary to obtain the gene sequence of species first, and then select an appropriate evolutionary model, and reliable calibration, such as the determining age of fossils (Ho and Duchene 2014; Ho and Phillips 2009). To clarify the phylogenetic position, divergence time and reasons of *S. aluco* from China, this study determined the complete mitochondrial genome of *S. aluco*, and used the mitochondrial genome combined with the mitochondrial genome of other birds in Strigiformes, The phylogenetic tree of Strigiformes was reconstructed. Fossil data are usually used to evaluate the divergence time of birds, and the divergence time of the Surniinae fossil is used as the correction point to analyze the divergence time of *Strix*, and the possible reasons for its divergence are fully discussed.

## MATERIAL AND METHOD

### Sample origin and DNA extraction

Part of muscle tissue was extracted from the leg of a *S. aluco* that died of unknown cause in the Rescue Center of Leigong Mountain National Nature Reserve, Qiandongnan Prefecture, Guizhou Province (26°49 ‘26.40 “N, 104°43’ 33.60” E). Stored in refrigerated boxes with built-in thermometers, keeping the temperature near freezing. Transported back to the lab for DNA extraction.

To extract DNA, we used the standardization CTAB method(Pinar et al., 2010; Lutz et al., 2011). And the DNA Sample Prep Kit was used to construct genomic DNA libraries.

### Sequencing and assembly

The Whole Genome Shotgun (WGS) strategy was used to construct the library (Roe, 2004). The Next Generation Sequencing (NGS) technology was used for paired-end sequencing(PE), based on the Illumina NovaSeq sequencing platform. The concentration and purity of DNA extracted from the samples were detected by Thermo Scientific NanoDrop 2000, and the integrity was detected by agarose electrophoresis and Agilent 2100 Bioanalyzer. Using the Covairs machine to break up DNA and fragment it. The gene library was constructed according to the shotgun method of Roe (2004). Agilent 2100 Bioanalyzer was used to detect the size of the library, and fluorescence quantitative detection was used to detect the total concentration of the library. The optimal amount of the library was selected and sequenced on Illumina. A single-stranded library was used as the template for bridge PCR amplification, and sequencing was performed while the synthesis.

After DNA extraction, purification, library construction and sequencing, the Raw image file obtained by sequencing is the first, and the Raw Data that can be read in FASTQ format is generated after multi-step transformation, that is, the offline data. The data transformation work is automatically completed by the sequencing platform. According to the statistics of Raw data, 7,947,240 Reads (each sequence read is called one read) were obtained, the total number of bases was 1192,086000bp, the percentage of fuzzy bases (uncertain bases) was 0.0016%, and the GC content was 44.58%. And base recognition accuracy of more than 99% accounted for 95.61% and base recognition accuracy of more than 99.9% accounted for 90.44%. The quality of the off-machine data should be tested through quality control, and the software used is FastQC(http://www.bioinformatics.babraham.ac.uk/projects/fastqc).

Sequencing data contains some low-quality reads with connectors, which will cause great interference to subsequent information analysis. In order to ensure the quality of subsequent information analysis, Fastp software (version 0.20.0) is needed to remove the contamination of sequencing connectors at the 3’ end. Low-quality sequences (sequences with an average Q value less than 20 and sequences with sequence length less than 50bp) were removed; The number of high-quality reads obtained was 7611,984, accounting for 95.78% of the raw data, and the number of bases of high-quality reads was 1123739765 bp, accounting for 94.27% of the raw data (Chen et al., 2018).

A5-miseq v20150522 (Coil et al., 2014) and SPAdesv3.9.0 (Bankevich et al., 2012) were used for the de novo assembly of high-quality next-generation sequencing data. Construct contig and scaffold sequences. The sequences were extracted according to the sequencing depth of de novo splicing sequences, and the sequences with high sequencing depth were compared with the nt library on NCBI by blastn (BLAST v2.2.31+), and the mitochondrial sequences of each splicing result were selected. Integration of splicing results: The mitochondrial splicing results obtained by different software above were combined with reference sequences, and collinearity analysis was performed using mummer v3.1 (Kurtz et al., 2004) software to determine the position relationship between contigs and fill gaps between contigs. The results were corrected using pilon v1.18 (Walker et al., 2014) software to obtain the final mitochondrial sequence. The complete mitochondrial genome sequence obtained by splicing was uploaded to the MITOS web server (http://mitos2.bioinf.uni-leipzig.de/index.py) for functional annotation (Bernt M et al., 2013). Among them, RefSeq 81 Metazoa is selected for Reference, The Genetic Code is set to a second set of vertebrate codons, and the rest are set according to the default parameters set by MITOS.

Through the above methods, the base composition of the whole mitochondrial genome, protein-coding genes, and rRNA genes was obtained. CGview visualization software was used to draw the mitochondrial complete genome circle map (Stothard et al.,2011).

### Mitochondrial genome data collection in Strigiformes

Currently, there are 30 species with mitochondrial genomes greater than 10000bp in GenBank, including 27 species of Strigidae, and 3 species of Tytonidae, All taxonomic classifications of the species follow the current version of the IOC WORLD BIRD LIST (12.2) (http://dx.doi.org/10.14344/IOC.ML.12.2), the registration number as shown in table 1.

**Table 1.**
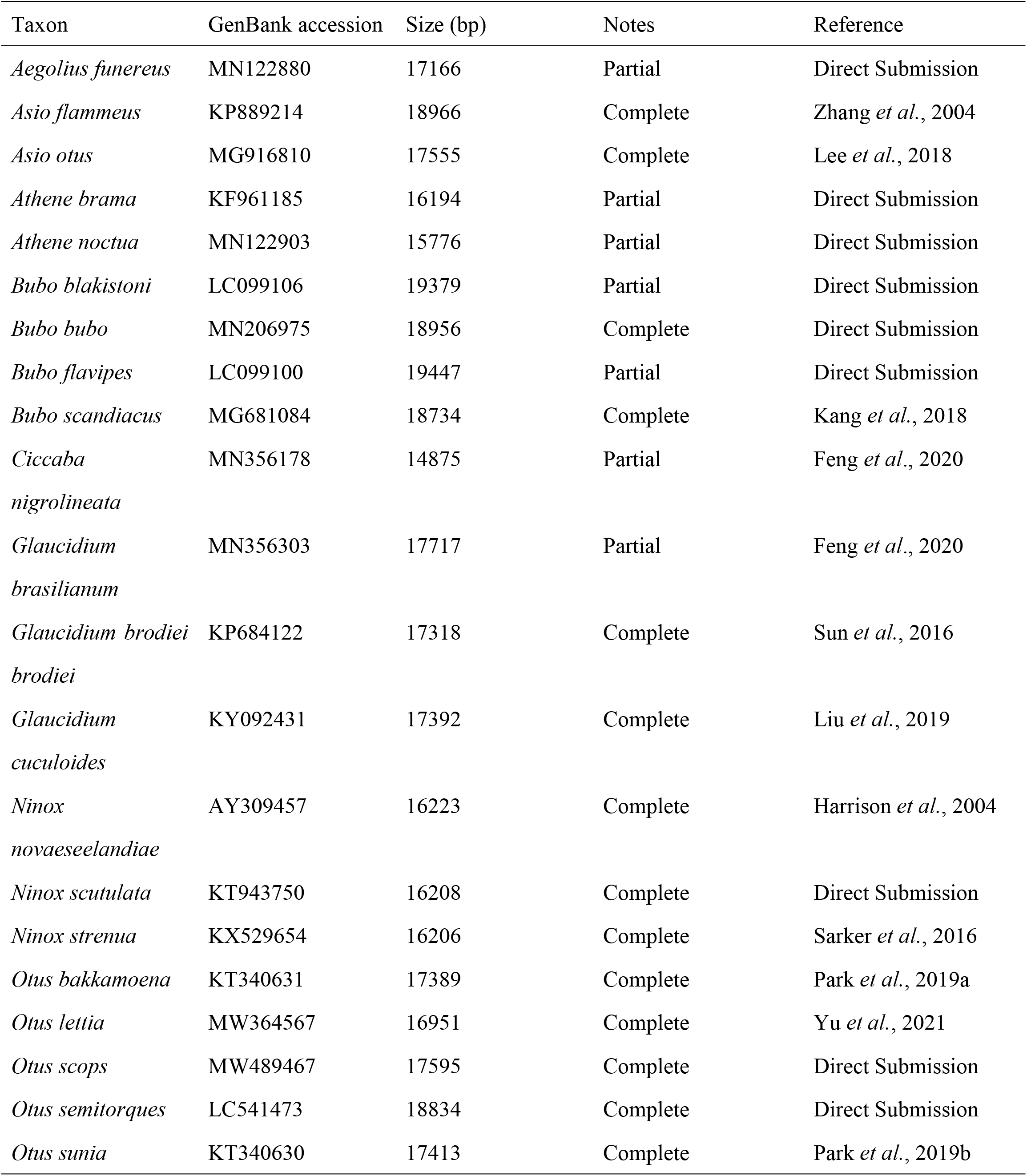
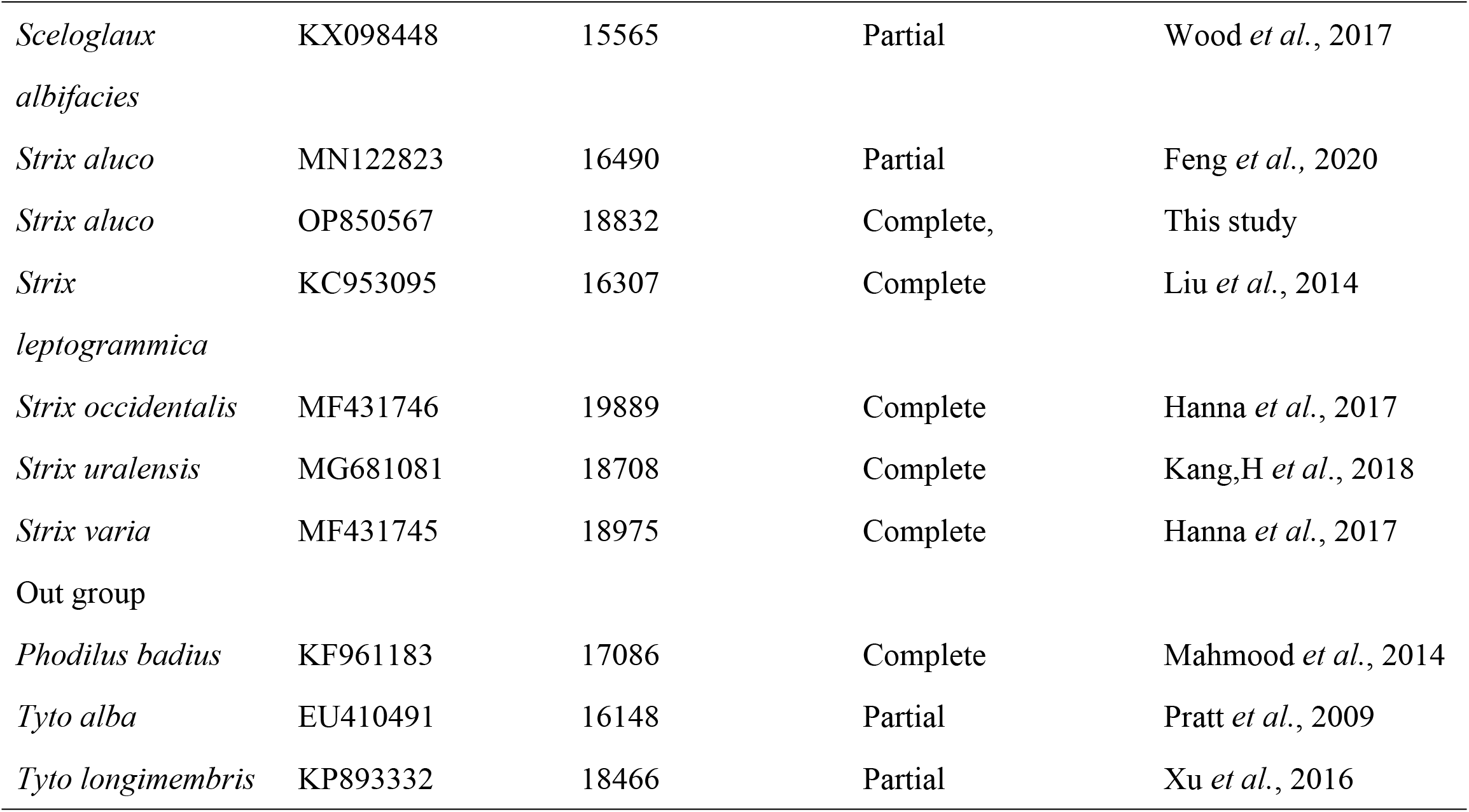
Mitochondrial genome sequences used in this study.

### Construction of phylogenetic trees

Using PhyloSuite software (download from: https://github.com/dongzhang0725/PhyloSuite/releases) (Zhang et al., 2020), Drag the 30 GenBank format files downloaded from NCBI and the GenBank format files of *S. aluco* sequence obtained by this sequencing into the main interface.

First series of standardized operations, the choice for Mitogenome sequence types, and then export the annotation error tRNA file, upload the ARWEN website (http://130.235.244.92/ARWEN/) modify comments, will be the site of the modified comments copy paste to modify after box. Secondly, 13 PCGs and 24 RNAs need to be extracted. The second set of 2 Vertebrate mitochondrial codes is selected here, and the extracted 13 PCGs and 24 RNAs are imported into MAFFT for multiple sequence alignment. Select the 37 gene files exported by MAFFT and import them into ‘concatenate sequence’, use the ‘-auto’ strategy and ‘normal’ alignment mode, and click start. Select the concatenated completion file and open the PartitionFinder 2.0 (Lafear et al., 2016), performed a greedy search using the Bayesian, and calculated the optimal partitioning strategy and model selection. Using a separate GTR+G model for each data block.

Select the result file of PartitionFinder 2.0 and complete the ML method in IQ-tree mode (Minh et al. 2020). Set Phodilus badius, Tyto alba and Tyto longimembris as out-group. under Edge-linked partition style for 10,000 replicates of ultrafast bootstrap (Hoang et al. 2018) et al. 2012), also select the result folder of PartitionFinder 2.0, open Mrbayes, set the out-groups, define parameters as Partition Models, run algebra as 2 parallel runs, 4 chains, 2,000,000 generations (must ensure the average standard deviation of split frequencies values were below 0.01), sampling freq is one sampling run for 1000 times, a burn-in number of initial 25% were burned.

### Divergence time evaluation

*Miosurnia diurna* fossils provide an approximate date of the origin of Surniinae. the age of the fossil-bearing sediments of the M. diurna is 6.0-9.5 Ma (Li et al., 2022), Surniinae may be composed of the species of Surnia, *Athene*, *Ninox*, and *Glaucidium* (Wink M et al., 2009), *M. diurna* fossil features are closer to the clade of Surnia +*Glaucidium*, Therefore, the origin times of *Glaucidium brasilianum, Glaucidium brodiei brodiei*, and *Glaucidium cuculoides* were set at 6Ma and 9.5Ma. The ‘NEX’ file obtained by concatenating 37 genes using the ‘concatenate sequence’ function in PhyloSuite was imported into BEAUti 2.6.7, (http://www.beast2.org/), Hasegawa-Kishino-Yano (HKY) model, with four gamma categories, Strict clock with 1.0 Clock rate, and with a Yule process (speciation) prior. Choose the *“Glaucidium brasilianum*, *Glaucidium brodiei brodiei, Glaucidium cuculoides”* (Sequence file name) to add Prior, Check the “monophyletic” option, set Mean to 6.0 /9.5 and Sigma to 0.1. A Markov chain Monte Carlo (MCMC) Bayesian analysis with a chain length of 10 000 000, and with states recorded every 1000 iterations, save to run using BEAST 2.6.7. Log files were assessed using TRACER 1.7.2 (http://tree.bio.ed.ac.uk/software/tracer/) to ensure posteriors were normally distributed and all statistics had attained effective sample sizes of >200, if ESS<200, try to optimize by adding 5000000 iterations (chain length) each time. A burn-in of 10% was discarded, and a maximum clade credibility tree was determined and Mean heights were chosen using TreeAnnotator 2.6.7. Finally, FigTree 1.4.4 was used to check the divergence time. Finally, use Adobe Illustrator 1.0.0.2 for visual editing.

## RESULTS

### Genome annotation

The total length of the mitochondrial genome sequence was 18,632 bp (Gen Bank entry number: OP850567). The genome annotation results showed that the total number of genes was 39, including 13 protein-coding genes, 22 tRNA genes, 2 rRNA genes, 2 *O_H_* genes, and 0 *O_L_* genes. Among them, 8 tRNA genes (trn-Q, trn-A, trn-N, trn-C, trn-Y, trn-P, trn-E, and trn-S2), one PCGs gene: nad6, are on the main chain (J chain); and the remaining 14 tRNA genes are trn-F, trn-V, trn-L2, trn-I, trn-M, trn-W, trn-D, trn-K, trn-G, trn-R, trn-H, trn-S1, trn-L1, trn-T; Two rRNA genes: rrn-S, rrn-L; And 12 PCGs genes encoding: nad1, nad2, nad3, nad4, nad4L, nad5, atp6, atp8, cox1, cox2, cox3, cytb on the secondary (N) chain (Table 2). There was no gene rearrangement (Fig. 1). The specific annotation results of each gene are shown in Table 2.

**Figure 1:**
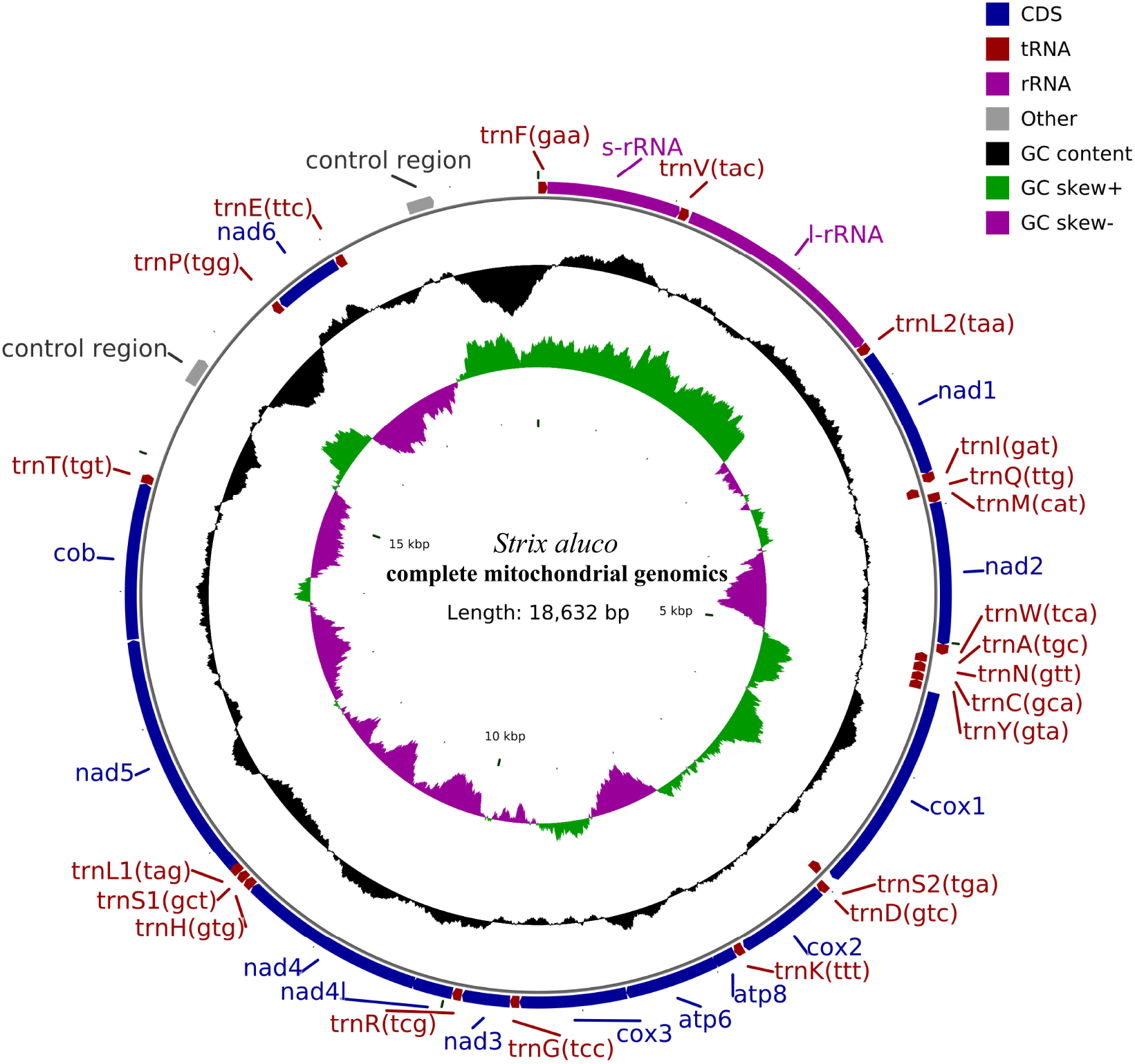
Complete mitochondrial genome of *S. aluco*. The total length of the mitochondrial genome of *S. aluco* is 18632Bp. The genes located on the N strand or J strand are positioned inside or outside the circle. Contains two D-Loop regions. The GC Skew+ region contains more Guanine than Cytosine, and the GC Skew-region contains more Cytosine than Guanine.

**Table 2.** Analysis of mitochondrial genome feature

| Feature | Strand | Position | Length<br>(bp) | Initiation<br>Codon | Stop<br>Codon | Anticodon | Intergenic<br>Nucleotide |
| --- | --- | --- | --- | --- | --- | --- | --- |
| trnF | N | 1-68 | 68 |  |  | GAA | -1 |
| rrnS | N | 68-1050 | 983 |  |  |  | -1 |
| trnV | N | 1050-1121 | 72 |  |  | TAC | 12 |
| rrnL | N | 1134-2709 | 1576 |  |  |  | -1 |
| trnL2 | N | 2709-2783 | 75 |  |  | TAA | 14 |
| nad1 | N | 2798-3757 | 960 | ATG | AGG |  | -2 |
| trnI | N | 3756-3827 | 72 |  |  | GAT | 11 |
| trnQ | J | 3839-3909 | 71 |  |  | TTG | -1 |
| trnM | N | 3909-3977 | 69 |  |  | CAT |  |
| nad2 | N | 3978-5018 | 1041 | ATG | TAG |  | -2 |
| trnW | N | 5017-5091 | 75 |  |  | TCA | 1 |
| trnA | J | 5093-5161 | 69 |  |  | TGC | 1 |
| trnN | J | 5163-5236 | 74 |  |  | GTT | 2 |
| trnC | J | 5239-5305 | 67 |  |  | GCA | -1 |
| trnY | J | 5305-5376 | 72 |  |  | GTA | 1 |
| cox1 | N | 5378-6928 | 1551 | ATG | AGG |  | -9 |
| trnS2 | J | 6920-6991 | 72 |  |  | TGA | 3 |
| trnD | N | 6995-7063 | 69 |  |  | GTC | 2 |
| cox2 | N | 7066-7749 | 684 | ATG | TAA |  | 9 |
| trnK | N | 7759-7828 | 70 |  |  | TTT | 1 |
| atp8 | N | 7830-7997 | 168 | ATG | TAA |  | -10 |
| atp6 | N | 7988-8671 | 684 | ATG | TAA |  | -1 |
| cox3 | N | 8671-9454 | 784 | ATG | T(AA) |  |  |
| trnG | N | 9455-9523 | 69 |  |  | TCC |  |
| nad3 | N | 9524-9875 | 352 | ATG | TAA |  | 2 |
| trnR | N | 9878-9946 | 69 |  |  | TCG | 1 |
| nad4l | N | 9948-10244 | 297 | ATG | TAA |  | -7 |
| nad4 | N | 10238-11615 | 1378 | ATG | T(AA) |  |  |
| trnH | N | 11616-11685 | 70 |  |  | GTG | 2 |
| trnS1 | N | 11688-11753 | 66 |  |  | GCT | 2 |
| trnL1 | N | 11756-11826 | 71 |  |  | TAG | -15 |
| nad5 | N | 11812-13647 | 1836 | ATG | TAA |  | 5 |
| cob | N | 13653-14795 | 1143 | ATG | TAA |  | 1 |
| trnT | N | 14797-14866 | 70 |  |  | TGT | 728 |
| OH_0 | N | 15595-15792 | 198 |  |  |  | 616 |
| trnP | J | 16409-16478 | 70 |  |  | TGG | 6 |
| nad6 | J | 16485-17006 | 522 | ATG | TAG |  | 3 |
| trnE | J | 17010-17083 | 74 |  |  | TTC | 584 |
| OH_1 | N | 17668-17865 | 198 |  |  |  | 766 |

### Phylogenetic analysis

In this study, both ML-tree and BI-tree show the same tree topology with good support, and it can be seen from the tree that Strigidae and Tytonidae are two distinct lineages under the owl shape. *Ciccaba nigrolineata* is nested in the Strix, and *Sceloglaux albifacies* is nested in the genus *Ninox. S. aluco* in this study is a sister group of *S. uralensis* (Kang et al., 2018), *Strix aluco* MN122823+ (*Strix aluco* OP850567+*Strix uralensis*) was formed with *S. aluco; Athene noctua* is a sister group of *Athene brama, Aegolius funereus* is a sister group of *Glaucidium cuculoides, Glaucidium brasilianum; Glaucidium brodiei brodiei, G. cuculoides, G. brasilianum, Athene noctua*, *A. brama*, *A. funereus* constitute the same group; *Strix* and *Bubo* are A sister group. And it forms an *Asio*+ (*Strix*+*Bubo*) monophyletic group with *Asio;* And a higher monophyletic group with *Otus*+[*Asio*+ (*Strix*+*Bubo*)], this monophyly simultaneously with (Sceloglaux albifacies+*Ninox*) monophyly exhibited as dyadic taxa.(Fig.2)

**Figure 2:**
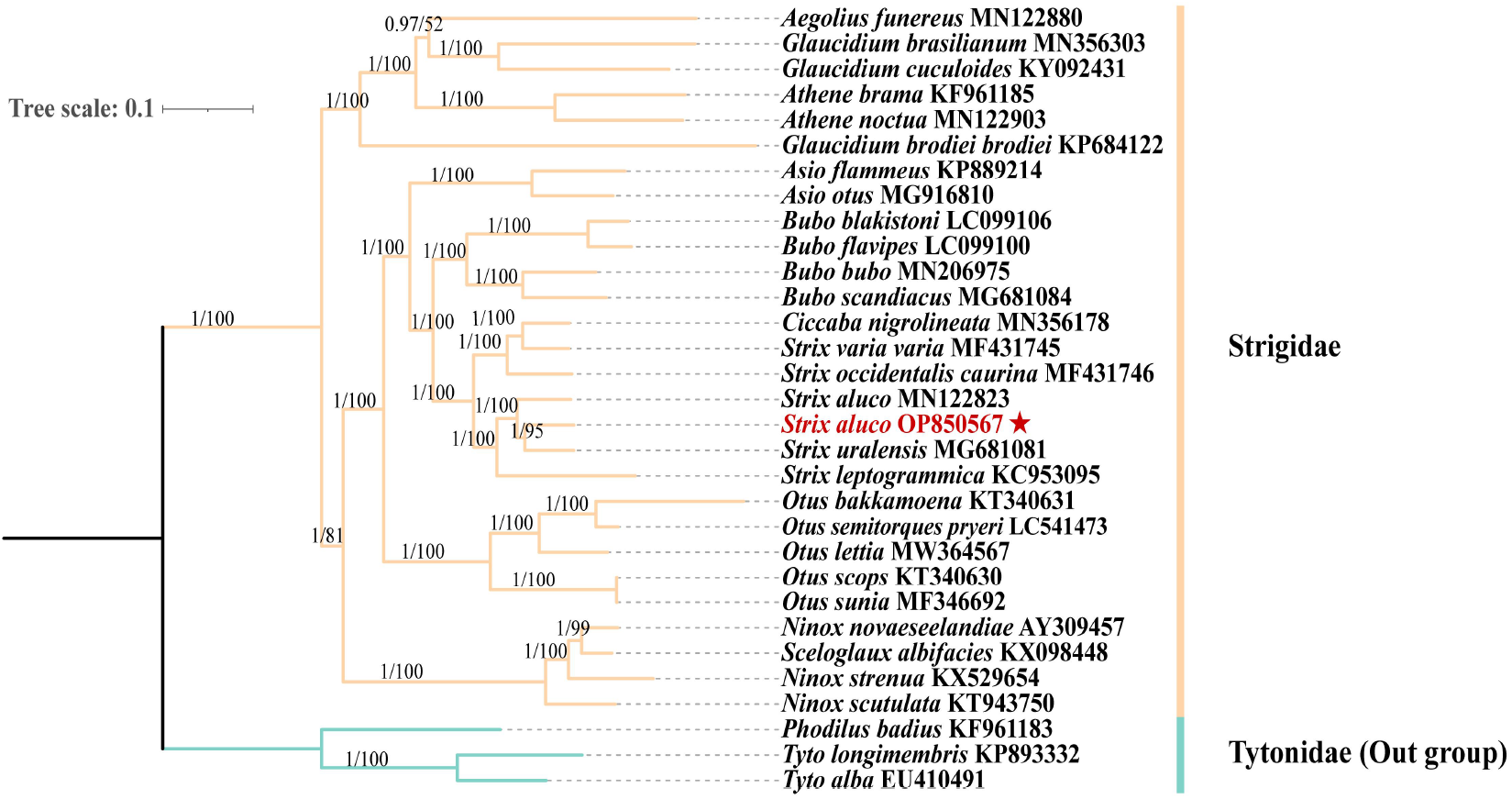
BI/ML--tree --. Bayesian phylogenetic tree of 37 genes (24 rRNAs, 13PCGs) from 31 species of Strigiformes. The node labels are BI/ML posterior probability and bootstrap support value respectively, and the scale indicates the probability of nucleotide change within each branch length. The GenBank of the sequences has been indicated next to the species name. Branches of different subfamilies are distinguished by different colors, with Tytoninae (with *Phodilus badius, Tyto longimembris* and *Tyto alba*) being the out-group. The *S. aluco* mitochondrial genome obtained by this sequencing has been marked by 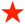.

### Divergence time evaluation

The divergence time tree based on 37 genomes shows that the time interval between Strigidae and Tytonidae from the common ancestor of Strigiformes was 8.69~13.76Ma. However, in the out-group, *Tyto alba, Tyto longimembris* and *Phodilus badius* diverged from the common ancestor about 4.57~7.24Ma. The divergence of Strigidae began about 6.81-10.79 Ma, and the earliest divergence of Surniinae occurred in Strigidae, and *Aegolius*, *Glaucidium*, and *Athene* occurred about 6.03-9.55 Ma. In this study, we found the divergence time between *S. aluco*(OP850567) and *S. aluco* of Margaryan. A (MN122823) was about 1.59~2.51Ma, and the divergence time between *S. aluco* and *S. uralensis* in China was about 1.38~2.19Ma. (Fig.3)

**Figure 3:**
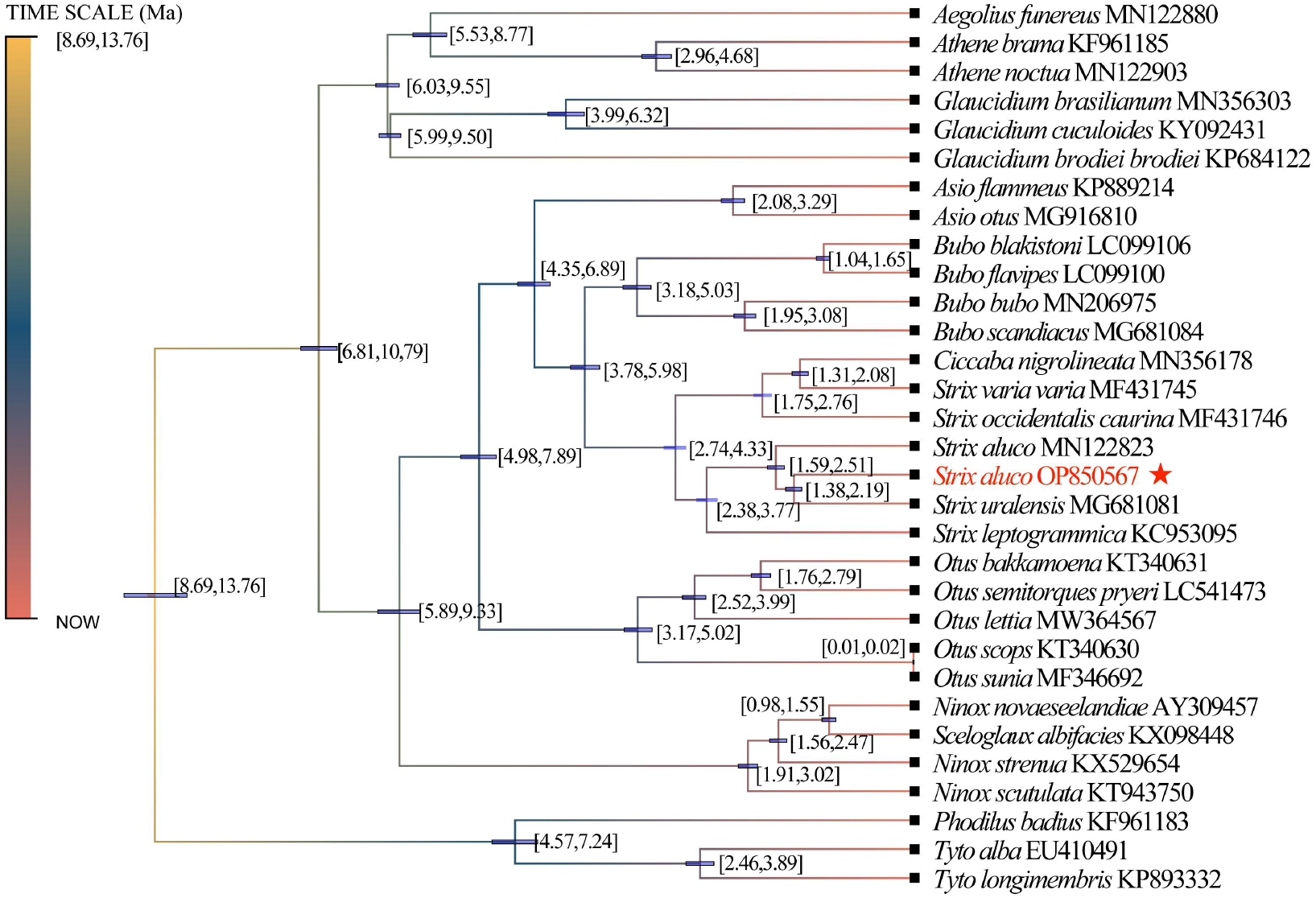
Divergence time tree. Through the divergence time tree obtained by Beast2.6.7 based on the Bayesian method, the node horizontal bar indicates that the posterior probability of this age interval is 95%, and the divergence time has been marked at the node.

## DISCUSSION

The mitochondrial genome structure of birds is a covalent double-chain loop structure, with a total of 37 genes, including 22 tRNAs, 2 rRNAs, 13 protein-coding genes (PCGs) and 1-2 non-coding control regions (D-loop). Where nad6 and 8 tRNA encoding genes (trnQ, trnA, trnN, trnC, trnY, trnS2, trnP and trnE) are on the J chain (light chain), The remaining 14 tRNAs, 2 rRNAs, 12 protein-coding genes and 1-2 non-coding control regions (D-loop) are all on the N chain (heavy chain) (Wolstenholme, 1992; Boore, 1999), and the complete mitochondrial genome structure of all birds was consistent (Hanna et al., 2017). The complete mitochondrial genome sequence of *S. aluco* obtained in this study was circular, with A total length of 18,632 bp and a GC content of 46.76%. Its composition was as follows: the proportion of Adenine bases in the total base column (A%) was 29.57%; The ratio of Guanine base to total base (G%) was 14.09%. The ratio of Cytosine base to total base (C%) was 32.67%. The ratio of Thymidine to the total base column (T%) was 23.66%. The start codon of all 13 PCGs is ATG, and the transcription stop codon is AGG, TAG and TAA. The content of A+T (53.23%) was higher than that of G+C (46.76%), which was consistent with the AT tendency of base bias in the vertebrate mitochondrial genome (Broughton et al., 2001; Ma et al., 2015), which is also consistent with the mitochondrial genome of other owls in Strigidae (Sun et al., 2020; Kang et al., 2018).

### Phylogenetic analysis of *S. aluco*

The BI and the ML tree have a consistent topology and each node has a high posterior probability. The phylogenetic tree of Strigiformes obtained by the mitochondrial genome in this study is consistent with the phylogenetic tree obtained by Li et al. (2022) through morphology. Our phylogenetic tree shows that Wink et al. (2009) compared Surnini (with Surnia, *Glaucidium* and *Taenioglaux*) Athenini (with *Athene*) and Aegolini (with *Aegolius*) under the Surniinae are feasible. In the phylogenetic tree constructed by Yu JJ et al. (2021), *C. nigrolineata* was also nested in *Strix*. *S. albifacies* has been largely extinct on the island of New Zealand, Wood JR et al. (2017) extracted its mitochondrial genome from museum specimens and suggested changing its name to *Ninox albifacies* because it has the same morphological structure and phylogenetic position as the genus *Ninox*. *S. aluco* in this study is a sister group of *S. uralensis* uploaded to GenBank by Kang et al. (2018). While S. aluco uploaded with Margaryan.A forms a monophyly of *Strix aluco* MN122823+ (*Strix aluco* OP850567+*Strix uralensis*). Compared with the mitochondrial genome of *S. aluco* obtained by Margaryan. A, *S. aluco* in China is more closely related to the *S. uralensis*, probably because at the beginning of the Pleistocene, the common ancestor of *S. aluco* MN122823 and *S. aluco* OP850567 had already been geographically isolated. The isolation of the Pleistocene refugium led to the divergence of the whole genome of the common ancestor of the forest owl at home and abroad. Foreign studies have shown that the Quaternary Period is characterized by a series of glacial-interglacial cycles (Woodruff, 2010), with the ancestors of modern species seeking refuge in a suitable environment. The existing species on the Qinghai-Tibet Plateau may be the result of rapid population expansion in relatively warm refugia during the Pleistocene glaciation and interglacial period, forming the current distribution pattern and genetic diversity (Gao et al., 2015). Legong Mountain in Guizhou Province just played the role of refugia for *S. aluco* during the Pleistocene glaciation period. Mitochondrial phylogeographic studies (Brito, 2005) show that the origin of *S. aluco* in Western Europe supports the “glacial refuge hypothesis”, and that the species survived in three allopatric refugees in the Iberian Peninsula, Italy and the Balkans, becoming the main source of *S. aluco* in Europe during the late glacial period. DNA barcoding technology also proved that the geographical barrier of the Strait of Gibraltar played an extremely important role in the phylogenetic history of *S. aluco* (Dona et al., 2016). The phylogenetic relationship of Strigidae forms a phylogenetic relationship of Surniinae+[*Ninox*+ (*Otus+ (Asio+* (*Strix*+*Bubo*)))], The conclusion of *Otus*+ (*Asio*+ (*Strix*+*Bubo*)) is consistent with the conclusion of Kang et al. (2018). According to the genome analysis of Strigidae birds in Madagascar, *Strix* is most closely related to *Bubo*, followed by *Otus* (Fuchs et al., 2008).

### Divergence time evaluation of *S. aluco*

On the Qinghai-Tibet Plateau, the impact of mountain uplift on the formation of modern species (<2.0 Ma) is limited, and researchers are more willing to believe that climate fluctuations played a key role in the formation of species in the Middle Pleistocene (Wang et al., 2018; Renner, 2016). During the Quaternary Period and Pleistocene (1.6~2.7Ma), there were severe climate shocks (Lisiecki and Raymo, 2005), which played a positive role in promoting the formation of species (Rull, 2008, 2011, 2015). The Pleistocene began 2.58 million years ago (2.58 Ma). Climate fluctuations during this period, especially during the ice age, affected the distribution of forests in the Northern Hemisphere and the evolution of species living in forests (Song et al., 2021). This series of climate fluctuations in the Pleistocene promoted species variation. This has led to species differentiation (Leonard et al., 2015). Glaciation has played an important role in influencing the population size, species and community genetic structure of today’s species (Svendsen et al., 2004; Hewitt 2000, 2004), the glacial-interglacial gyrations of the same period also affected the distribution of species (Kozma et al., 2018; Zhao et al., 2013; Hung et al., 2014; Mays Jr et al., 2018), Glacial-interglacial cycles led to periodic shifts in glacial refuges for Pleistocene birds (Nadachowska-Brzyska et al., 2015), the isolation of glacier refugia will lead to the divergence of the whole genome of species, thus forming different species (Provost et al., 2022), this should be the reason why the common ancestor of *S. aluco* MN122823 and *S. aluco* OP850567 (this study) diverged at 1.59~2.51Ma, genetic divergence of the same lineage due to the isolation of refugees leads to lineage divergence. All kinds of species generally begin to migrate to the best habitat during the warm climate period (Claramunt and Cracraft, 2015), in particular, species adapted to low altitudes in the early stage of climate change will move to high altitudes at this time, resulting in the reproductive isolation of species in the two places (Wiens, 2004). During the Pleistocene-Holocene (1.10 ~ 0.60 Ma), the Qinghai-Tibet Plateau experienced three stages of rapid uplift, with mountains forming, climate-changing from moist and warm to dry and cold, and forests retreating to the edge of the plateau (Wang et al., 2008). The Quaternary Period climate shock led to the initial formation of the existing forest and mountain distribution pattern in the Northern Hemisphere. Birds began to be widely distributed after leaving the glacier refuge at the end of the glacier and initially formed the existing distribution pattern (Pujolar et al., 2022). The long-tailed forest owl may have moved north at this time and thus diverged from *S. aluco*. In addition, the rapid uplift of the Qinling Mountains from the end of the Early Pleistocene to the Middle Pleistocene may have made the Qinling Mountains a barrier to north-south bird communication (Li et al., 2019). The rapid uplift of the Qinling Mountains prevented communication between *S. aluco* and the common ancestor of *S.uralensis*, which was originally distributed on the north and south sides.

In summary, by sequencing the complete mitochondrial genome of *S. aluco*, and mapping its phylogenetic tree and divergence time tree, the phylogenetic relationship of Strigiformes (Tytoninae + Phodilinae) + (Striginae+Ninoxinae+Surniinae) is summarized, with Tytonidae including Tytoninae (with *Tyto*) and Phodilinae (with *Phodilus*) as the out-group, Strigidae comprises Striginae (with *Asio, Bubo, Strix, Ciccaba* and *Otus*) +Ninoxinae+Surniinae (with *Athenini*, *Aegolini* and *Glaucidium*). The divergence time tree showed that the divergence time between *S. aluco* of China and *S. aluco* of other countries was about 1.59~2.51Ma, suggesting that the common ancestor of *S. aluco* at home and abroad was separated by geographical isolation at the beginning of the Pleistocene. The divergence between *S. aluco* and *S. uralensis* in China was about 1.38~2.19Ma. During this time, the rapid uplift of the Qinling Mountains led to the divergence of the ancestors of *Strix* on the north and south sides of the Chinese mainland. At the same time, due toClimatic oscillation in the Pleistocene, the existing *S. aluco* population on the Qinghai-Tibet Plateau may have rapidly expanded in relatively warm shelters such as Leigong Mountain to form the current distribution pattern.

## Acknowledge

Thank Rescue Center of Leigong Mountain National Nature Reserve, Qiandongnan Prefecture, Guizhou Province (26° 49 ‘26.40 “N, 104° 43’ 33.60” E) for provides the genes of strix aluco

## Author contributions

Conceptualization: Y, W., H, Z., Y, Z., Z, L.; Methodology: Y, W., H, Z.; Software: H, Z.; Formal analysis: H, Z.; Investigation:Y, W., Y, X., Y, Z., Z, L., B, L., X, L.; Resources: Rescue Center of Leigong Mountain National Nature Reserve, Qiandongnan Prefecture, Guizhou Province; Data curation: Z, H.; Writing - original draft:Y, W., Z, H.; Writing - review & editing: Y, Z., Z, L. Y, X.; Visualization: H, Z.,Y, W.; Supervision: Y, X.; Project administration: Y, X.; Funding acquisition: Y, X.; Y, W.

## Data ability

The complete mitochondrial genome of *S.aluco* has been uploaded to NCBI, GenBank accession number: OP850567.

## REFERENCE

Bankevich, A., Nurk, S., Antipov, D., Gurevich, A. A., Dvorkin, M., Kulikov, A. S., Lesin, V. M., Nikolenko, S. I., Pham, S., Pevzner, P. A. et al. (2012). SPAdes: a new genome assembly algorithm and its applications to single-cell sequencing. Journal of computational biology, 19, 455–477. doi:10.1089/cmb.2012.0021

Bernt, M., Donath, A., Jühling, F., Externbrink, F., Florentz, C., Fritzsch, G., Pütz, J., Middendorf, M. and Stadler, P. F. (2013). MITOS: improved de novo metazoan mitochondrial genome annotation. Molecular phylogenetics and evolution, 69, 313–319. doi:10.1016/j.ympev.2012.08.023

Boore, J. L. (1999). Animal mitochondrial genomes. Nucleic acids research, 27, 1767–1780.

Brito, P. H. (2005). The influence of Pleistocene glacial refugia on tawny owl genetic diversity and phylogeography in western Europe. Molecular Ecology, 14, 3077–3094. doi:10.1111/J.1365-294X.2005.02663.X

Broughton, R. E., Milam, J. E. and Roe, B. A. (2001). The complete sequence of the zebrafish (Danio rerio) mitochondrial genome and evolutionary patterns in vertebrate mitochondrial DNA. Genome research, 11, 1958–1967. doi:10.1101/gr.156801

Chen, S., Zhou, Y., Chen, Y. and Gu, J. (2018). fastp: an ultra-fast all-in-one FASTQ preprocessor. Bioinformatics, 34, i884–i890. doi:10.1101/274100

Claramunt, S. and Cracraft, J. (2015). A new time tree reveals Earth history’s imprint on the evolution of modern birds. Science advances, 1, e1501005. doi:10.1126/sciadv.1501005

Coil, D., Jospin, G. and Darling, A. E. (2015). A5-miseq: an updated pipeline to assemble microbial genomes from Illumina MiSeq data. Bioinformatics, 31, 587–589. doi:10.1093/bioinformatics/btu661

Comay, O., Ezov, E., Yom-Tov, Y. and Dayan, T. (2022). In Its Southern Edge of Distribution, the Tawny Owl (Strix aluco) Is More Sensitive to Extreme Temperatures Than to Rural Development. Animals, 12, 641. doi:10.3390/ani12050641

Doña, J., Ruiz-Ruano, F. J. and Jovani, R. (2016). DNA barcoding of Iberian Peninsula and North Africa Tawny Owls Strix aluco suggests the Strait of Gibraltar as an important barrier for phylogeography. Mitochondrial DNA Part A, 27, 4475–4478. doi:10.3109/19401736.2015.1089573

Drovetski, S. V. (2003). Plio-Pleistocene climatic oscilations, Holarctic biogeography and speciation in an avian subfamily. Journal of Biogeography, 30, 1173–1181. doi:10.1046/j.1365-2699.2003.00920.x

Feng, S., Stiller, J., Deng, Y., Armstrong, J., Fang, Q. I., Reeve, A. H., Xie, D., Faircloth, B. C., Petersen, B., Zhang, G. et al. (2020). Dense sampling of bird diversity increases power of comparative genomics. Nature, 587, 252–257. doi:10.1038/s41586-021-03473-8

Field, D. J., Bercovici, A., Berv, J. S., Dunn, R., Fastovsky, D. E., Lyson, T. R., Vajda, V. and Gauthier, J. A. (2018). Early evolution of modern birds structured by global forest collapse at the end-Cretaceous mass extinction. Current Biology, 28, 1825–1831. doi:10.1016/j.cub.2018.04.062

Fuchs, J., Pons, J. M., Goodman, S. M., Bretagnolle, V., Melo, M., Bowie, R. C., Currie, D., Safford, R., Virani, M. Z., Thomsett, S. et al.(2008). Tracing the colonization history of the Indian Ocean scops-owls (Strigiformes: Otus) with further insight into the spatio-temporal origin of the Malagasy avifauna. BMC Evolutionary Biology, 8, 1–15. doi:10.1186/1471-2148-8-197

Gao, Y. D., Zhang, Y., Gao, X. F. and Zhu, Z. M. (2015). Pleistocene glaciations, demographic expansion and subsequent isolation promoted morphological heterogeneity: A phylogeographic study of the alpine Rosa sericea complex (Rosaceae). Scientific Reports, 5, 1–15. doi:10.1038/srep11698

Grantsmanship, E. E., Kuznetsov, A. N. and Panyutina, A. A. (2016) Kinematic constituents of the extreme head turn of Strix aluco estimated by means of CT-scanning. In Doklady Biological Sciences. Pleiades Publishing, 466: 24–27. doi:10.1134/S0012496616010087

Hanna, Z. R., Henderson, J. B., Sellas, A. B., Fuchs, J., Bowie, R. C. and Dumbacher, J. P. (2017). Complete mitochondrial genome sequences of the northern spotted owl (Strix occidentalis caurina) and the barred owl (Strix varia; Aves: Strigiformes: Strigidae) confirm the presence of a duplicated control region. PeerJ, 5, e3901. doi:10.7717/peerj.3901

Haring, E., Kruckenhauser, L., Gamauf, A., Riesing, M. J. and Pinsker, W. (2001). The complete sequence of the mitochondrial genome of Buteo buteo (Aves, Accipitridae) indicates an early split in the phylogeny of raptors. Molecular Biology and Evolution, 18, 1892–1904. doi:10.1093/oxfordjournals.molbev.a003730

Harrison, G. L., McLenachan, P. A., Phillips, M. J., Slack, K. E., Cooper, A. and Penny, D. (2004). Four new avian mitochondrial genomes help get to basic evolutionary questions in the late Cretaceous. Molecular Biology and Evolution, 21, 974–983. doi:10.1093/molbev/msh065

Heidrich, P. and Wink, M. (1994). Tawny owl (Strix aluco) and Hume’s Tawny owl (Strix butleri) are distinct species: evidence from nucleotide sequences of the cytochrome b gene. Zeitschrift für Naturforschung C, 49, 230–234.

Hewitt, G. (2000). The genetic legacy of the Quaternary ice ages. Nature, 405, 907–913. doi:10.1038/35016000

Hewitt, G. (2004) Genetic consequences of climatic oscillations in the Quaternary. Philos Trans R Soc B Biol Sci. 359:183–195. doi:10.1098/rstb.2003.1388

Ho, S. Y. and Phillips, M. J. (2009). Accounting for calibration uncertainty in phylogenetic estimation of evolutionary divergence times. Systematic biology, 58, 367–380. doi:10.1093/sysbio/syp035

Ho, S. Y. and Duchêne, S. (2014). Molecular-clock methods for estimating evolutionary rates and timescales. Molecular ecology, 23, 5947–5965. doi:10.1111/mec.12953

Holm, S. R. and Svenning, J. C. (2014). 180,000 years of climate change in Europe: avifaunal responses and vegetation implications. Plos one, 9, e94021. doi:10.1371/journal.pone.0094021

Hung, C. M., Shaner, P. J. L., Zink, R. M., Liu, W. C., Chu, T. C., Huang, W. S. and Li, S. H. (2014). Drastic population fluctuations explain the rapid extinction of the passenger pigeon. Proceedings of the National Academy of Sciences, 111, 10636–10641. doi:10.1073/pnas.1401526111

Kang, H., Li, B., Ma, X. and Xu, Y. (2018). Evolutionary progression of mitochondrial gene rearrangements and phylogenetic relationships in Strigidae (Strigiformes). Gene, 674, 8–14. doi:10.1016/j.gene.2018.06.066

Karell, P., Ahola, K., Karstinen, T., Zolei, A. and Brommer, J. E. (2009). Population dynamics in a cyclic environment: consequences of cyclic food abundance on tawny owl reproduction and survival. Journal of Animal Ecology, 78, 1050–1062. doi:10.1111/j.1365-2656.2009.01563.x

Koparde, P., Mehta, P., Reddy, S., Ramakrishnan, U., Mukherjee, S. and Robin, V. V. (2018). The critically endangered forest owlet Heteroglaux blewitti is nested within the currently recognized Athene clade: A century-old debate addressed. PloS one, 13, e0192359. doi:10.1371/journal.pone.0192359

Kozma, R., Lillie, M., Benito, B. M., Svenning, J. C. and Höglund, J. (2018). Past and potential future population dynamics of three grouse species using ecological and whole genome coalescent modeling. Ecology and evolution, 8, 6671–6681. doi:10.1002/ece3.4163

Ksepka, D. T. and Phillips, M. J. (2015). Avian Diversification Patterns across the K-Pg Boundary: Influence of Calibrations, Datasets, and Model Misspecification1. Annals of the Missouri Botanical Garden, 100, 300–328. doi:10.3417/2014032

Kurtz, S., Phillippy, A., Delcher, A. L., Smoot, M., Shumway, M., Antonescu, C. and Salzberg, S. L. (2004). Versatile and open software for comparing large genomes. Genome biology, 5, 1–9. doi:10.1186/gb-2004-5-2-r12

Lamb, A. M., Gonçalves da Silva, A., Joseph, L., Sunnucks, P. and Pavlova, A. (2019). Pleistocene-dated biogeographic barriers drove divergence within the Australo-Papuan region in a sex-specific manner: an example in a widespread Australian songbird. Heredity, 123, 608–621. doi:10.1038/s41437-019-0206-2

Leonard, J. A., den Tex, R-J., Hawkins, M. T. R., Munoz-Fuentes, V., Thorington, R. and Maldonado, J. E. (2015). Phylogeography of vertebrates on the Sunda Shelf: a multi-species comparison. Journal of Biogeography, 42, 871–879. doi:10.1111/jbi.12465

Lee, M. Y., Lee, S. M., Jeon, H. S., Lee, S. H., Park, J. Y. and An, J. (2018). Complete mitochondrial genome of the Northern Long-eared Owl (Asio otus Linnaeus, 1758) determined using next-generation sequencing. Mitochondrial DNA Part B, 3, 494–495. doi:10.1080/23802359.2018.1451260

Li, J., Song, G., Liu, N., Chang, Y. and Bao, X. (2019). Deep south-north genetic divergence in Godlewski’s bunting (Emberiza godlewskii) related to uplift of the Qinghai-Tibet Plateau and habitat preferences. BMC evolutionary biology, 19, 1–13. doi:10.1186/s12862-019-1487-z

Li, Z.H., Stidham, T. A., Zheng, X., Wang, Y., Zhao, T., Deng, T. and Zhou, Z. (2022). Early evolution of diurnal habits in owls (Aves, Strigiformes) documented by a new and exquisitely preserved Miocene owl fossil from China. Proceedings of the National Academy of Sciences, 119, e2119217119. doi:10.1073/pnas.2119217119

Liu, G., Zhou, L. and Gu, C. (2014). The complete mitochondrial genome of Brown wood owl Strix leptogrammica (Strigiformes: Strigidae). Mitochondrial DNA: The Journal of DNA Mapping. 25, 370–371. doi:10.3109/19401736.2013.803540

Liu, G., Zhou, L. and Zhao, G. (2019). Complete mitochondrial genomes of five raptors and implications for the phylogenetic relationships between owls and nightjars. PeerJ Preprints. No. e27478v1. doi:10.7287/peerj.preprints.27478v1

Lisiecki, L.E. and Raymo, M. E. (2005). A Pliocene-Pleistocene stack of 57 globally distributed benthic δ18O records. Paleoceanography 20, 1–17. doi:10.1029/2004pa001071

Lutz, K. A., Wang, W., Zdepski, A. and Michael, T. P. (2011). Isolation and analysis of high quality nuclear DNA with reduced organellar DNA for plant genome sequencing and resequencing. BMC biotechnology, 11, 1–9. doi:10.1186/1472-6750-11-54

Ma, Z., Yang, X., Bercsenyi, M., Wu, J., Yu, Y., Wei, K., et al. and Yang, R. (2015). Comparative mitogenomics of the genus Odontobutis (Perciformes: Gobioidei: Odontobutidae) revealed conserved gene rearrangement and high sequence variations. International journal of molecular sciences, 16, 25031–25049. doi:10.3390/ijms161025031

Mahmood, M. T., McLenachan, P. A., Gibb, G. C. and Penny, D. (2014). Phylogenetic position of avian nocturnal and diurnal raptors. Genome biology and evolution, 6, 326–332. doi:10.1093/gbe/evu016

Mays Jr, H. L., Hung, C. M., Shaner, P. J., Denvir, J., Justice, M., Yang, S. F., Roth, T. L., Fan, J., Rekulapally, S., Primerano, D. A. et al.(2018). Genomic analysis of demographic history and ecological niche modeling in the endangered Sumatran rhinoceros Dicerorhinus sumatrensis. Current Biology, 28, 70–76. doi:10.1016/j.cub.2017.11.021

Nadachowska-Brzyska, K., Li, C., Smeds, L., Zhang, G. and Ellegren, H. (2015). Temporal dynamics of avian populations during Pleistocene revealed by whole-genome sequences. Current Biology, 25, 1375–1380. doi:10.1016/j.cub.2015.03.047

Obuch, J. (2011). Spatial and temporal diversity of the diet of the tawny owl (Strix aluco). Raptor Journal, 5, 1–120. doi:10.2478/v10262-012-0057-8

Park, C. E., Kim, M. C., Ibal, J. C. P., Pham, H. Q., Park, H. C. and Shin, J. H. (2019a). The complete mitochondrial genome sequence of Otus bakkamoena (Aves, Strigiformes, Strigidae). Mitochondrial DNA Part B, 4, 775–776. doi:10.1080/23802359.2019.1565979

Park, C. E., Kim, M. C., Ibal, J. C. P., Pham, H. Q., Park, H. C. and Shin, J. H. (2019b). The complete mitochondrial genome sequence of Otus scops (Aves, Strigiformes, Strigidae). Mitochondrial DNA Part B, 4, 764–765. doi:10.1080/23802359.2019.1565973

Pratt, R. C., Gibb, G. C., Morgan-Richards, M., Phillips, M. J., Hendy, M. D. and Penny, D. (2009). Toward resolving deep Neoaves phylogeny: data, signal enhancement, and priors. Molecular Biology and Evolution, 26, 313–326. doi:10.1093/molbev/msn248

Provost, K., Shue, S. Y., Forcellati, M. and Smith, B. T. (2022). The genomic landscapes of desert birds form over multiple time scales. Molecular Biology and Evolution, 39, msac200. doi:10.1093/molbev/msac200

Pujolar, J. M., Blom, M. P. K., Reeve, A. H., Kennedy, J. D., Marki, P. Z., Korneliussen, T. S., Freeman, B.G., Sam, K., Linck, E., Haryoko, T. et al. (2022) The formation of avian montane diversity across barriers and along elevational gradients. Nat Commun 13, 268. doi:10.1038/s41467-021-27858-5

Pinar, A., Akyön, Y., Alp, A. and Ergüven, S. Ì. B. E. L. (2010). Adaptation of a sensitive DNA extraction method for detection of Entamoeba histolytica by real-time polymerase chain reaction. Mikrobiyoloji bulteni, 44, 453–459.

Renner, S.S. (2016) Available data point to a 4-km-high Tibetan Plateau by 40 Ma, but 100 molecular-clock papers have linked supposed recent uplift to young node ages. Journal of Biogeography, 43, 1479–1487. doi:10.1111/jbi.12755

Roe, B. A. (2004). Shotgun library construction for DNA sequencing. In Bacterial artificial chromosomes. Humana Press. 171–187.

Rull, V. (2008) Speciation timing and neotropical biodiversity: the Tertiary-Quaternary debate in the light of molecular phylogenetic evidence. Mol. Ecol. 17, 2722–2729. doi:10.1111/j.1365-294X.2008.03789.x

Rull, V. (2015). Pleistocene speciation is not refuge speciation. J. Biogeogr. 42, 602–604. doi:10.1111/jbi.12440

Rull, V. (2011).. Neotropical biodiversity: timing and potential drivers. Trends Ecol. Evol. 26, 508–513. doi:10.1016/j.tree.2011.05.011

Salter, J. F., Oliveros, C. H., Hosner, P. A., Manthey, J. D., Robbins, M. B., Moyle, R. G., Faircloth, B. C. and Brumfield, R. T. (2020). Extensive paraphyly in the typical owl family (Strigidae). The Auk, 137, ukz070. doi:10.1093/auk/ukz070

Sarker, S., Das, S., Forwood, J., Helbig, K. and Raidal, S. R. (2016). The complete mitochondrial genome sequence of an Endangered powerful owl (Ninox strenua). Mitochondrial DNA Part B, 1, 722–723. doi:10.1080/23802359.2016.1229588

Solonen, T. and Karhunen, J. (2002). Effects of variable feeding conditions on the Tawny Owl Strix aluco near the northern limit of its range. Ornis Fennica, 79, 121–131.

Song, K., Gao, B., Halvarsson, P., Fang, Y., Klaus, S., Jiang, Y. X., Swenson, J. E., Sun, Y. H. and Höglund, J. (2021). Demographic history and divergence of sibling grouse species inferred from whole genome sequencing reveal past effects of climate change. BMC ecology and evolution, 21, 1–10. doi:10.1186/s12862-021-01921-7

Stothard, P. and Wishart, D. S. (2005). Circular genome visualization and exploration using CGView. Bioinformatics, 21, 537–539. doi:10.1093/bioinformatics/bti054

Sun, X., Zhou, W., Sun, Z., Qian, L., Zhang, Y., Pan, T. and Zhang, B. (2016). The complete mitochondrial genome of Glaucidium brodiei (Strigiformes: Strigidae). Mitochondrial DNA Part A, 27, 2508–2509. doi:10.3109/19401736.2015.1036252

Sun, C. H., Liu, H. Y., Min, X. and Lu, C. H. (2020). Mitogenome of the little owl Athene noctua and phylogenetic analysis of Strigidae. International journal of biological macromolecules, 151, 924–931. doi:10.1016/j.ijbiomac.2020.02.238

Sunde, P., Bølstad, M. S. and Desfor, K. B. (2003). Diurnal exposure as a risk sensitive behaviour in tawny owls Strix aluco?. Journal of Avian Biology, 34, 409–418. doi:DOI:10.1111/j.0908-8857.2003.03105.x

Svendsen, J. I., Alexanderson, H., Astakhov, V. I., Demidov, I., Dowdeswell, J. A., Funder, S., Gataullin, V., Henriksen, M., Hjort, C., Houmark-Nielsen, M. et al. (2004). Late Quaternary ice sheet history of northern Eurasia. Quaternary Science Reviews, 23, 1229–1271. doi:10.1016/j.quascirev.2003.12.008

Tuinen, M. V., Sibley, C. G. and Hedges, S. B. (2000). The early history of modern birds inferred from DNA sequences of nuclear and mitochondrial ribosomal genes. Molecular Biology and Evolution, 17, 451–457.

Uva, V., Päckert, M., Cibois, A., Fumagalli, L. and Roulin, A. (2018). Comprehensive molecular phylogeny of barn owls and relatives (Family: Tytonidae and their six major Pleistocene radiations. Molecular phylogenetics and evolution, 125, 127–137. doi:10.1016/j.ympev.2018.03.013

Voelker, G. (2010). Repeated vicariance of Eurasian songbird lineages since the late Miocene. Journal of biogeography, 37, 1251–1261. doi:10.1111/j.1365-2699.2010.02313.x

Walker, B. J., Abeel, T., Shea, T., Priest, M., Abouelliel, A., Sakthikumar, S., Cuomo, C. A., Zeng, Q., Wortman, J., Young, S. K. et al.(2014). Pilon: an integrated tool for comprehensive microbial variant detection and genome assembly improvement. PloS one, 9, e112963. doi:10.1371/journal.pone.0112963

Wang, C., Zhao, X., Liu, Z., Lippert, P. C., Graham, S. A., Coe, R. S., Yi, H., Zhu, L., Liu, S. and Li, Y. (2008). Constraints on the early uplift history of the Tibetan Plateau. Proceedings of the National Academy of Sciences, 105, 4987–4992. doi:10.1073/pnas.0703595105

Wang, P., Yao, H., Gilbert, K. J., Lu, Q., Hao, Y., Zhang, Z. and Wang, N. (2018). Glaciation-based isolation contributed to speciation in a Palearctic alpine biodiversity hotspot: evidence from endemic species. Molecular phylogenetics and evolution, 129, 315–324. doi:10.1016/j.ympev.2018.09.006

Wiens, J. J. (2004). Speciation and ecology revisited: phylogenetic niche conservatism and the origin of species. Evolution, 58, 193–197. doi:10.1111/j.0014-3820.2004.tb01586.x

Wink, M. and Heidrich, P. (2000). Molecular systematics of owls (Strigiformes) based on DNA-sequences of the mitochondrial cytochrome b gene. Raptors at risk, 819–828. doi:?

Wink, M., El-Sayed, A. A., Sauer-Gürth, H. and Gonzalez, J. (2009). Molecular phylogeny of owls (Strigiformes) inferred from DNA sequences of the mitochondrial cytochrome b and the nuclear RAG-1 gene. Ardea, 97, 581–591. doi:10.5253/078.097.0425

Wolstenholme, D. R. (1992). Animal mitochondrial DNA: structure and evolution. International review of cytology, 141, 173–216.

Woodruff, D. S. (2010). Biogeography and conservation in Southeast Asia: how 2.7 million years of repeated environmental fluctuations affect today’s patterns and the future of the remaining refugial-phase biodiversity. Biodiversity and Conservation, 19, 919–941. doi:10.1007/s10531-010-9783-3

Wood, J. R., Mitchell, K. J., Scofield, R. P., De Pietri, V. L., Rawlence, N. J. and Cooper, A. (2017). Phylogenetic relationships and terrestrial adaptations of the extinct laughing owl, Sceloglaux albifacies (Aves: Strigidae). Zoological Journal of the Linnean Society, 179, 907–918. doi:10.1111/zoj.12483

Xu, P., Li, Y., Miao, L., Xie, G. and Huang, Y. (2016). Complete mitochondrial genome of the Tyto longimembris (Strigiformes: Tytonidae). Mitochondrial DNA Part A, 27, 2481–2482. doi:10.3109/19401736.2015.1033708

Yan, C., Mou, B., Meng, Y., Tu, F., Fan, Z., Price, M., Zhang, X. and Zhang, X.(2017). A novel mitochondrial genome of Arborophila and new insight into Arborophila evolutionary history. Plos one, 12, e0181649. doi:10.1371/journal.pone.0181649

Yu, J., Liu, J., Li, C., Wu, W., Feng, F., Wang, Q., Ying, X., Qi, D. and Qi, G. (2021). Characterization of the complete mitochondrial genome of Otus lettia: exploring the mitochondrial evolution and phylogeny of owls (Strigiformes). Mitochondrial DNA Part B, 6, 3443–3451. doi:10.1080/23802359.2021.1995517

Zhao, S., Zheng, P., Dong, S., Zhan, X., Wu, Q., Guo, X., Hu, Y., He, W., Zhang, S. and Wei, F. (2013). Whole-genome sequencing of giant pandas provides insights into demographic history and local adaptation. Nature Genetics, 45, 67–U99. doi:10.1038/ng.2494

Zhang, Y., Song, T., Pan, T., Sun, X., Sun, Z., Qian, L. and Zhang, B. (2016). Complete sequence and gene organization of the mitochondrial genome of Asio flammeus (Strigiformes, strigidae). Mitochondrial DNA Part A, 27, 2665–2667. doi:10.3109/19401736.2015.1043538

Zhang, D., Gao, F., Jakovlić, I., Zou, H., Zhang, J., Li, W. X. and Wang, G. T. (2020). PhyloSuite: an integrated and scalable desktop platform for streamlined molecular sequence data management and evolutionary phylogenetics studies. Molecular ecology resources, 20, 348–355. doi:10.1111/1755-0998.13096

